## Supplementary material for "Predicting the Functional Impact of KCNQ1 Variants with Artificial Neural Networks": Figure S1 Figure S2 Figure S3 Figure S4 Figure S5 Figure S6 Figure S7 Figure S8 Table S1

### Supporting information

#### Biophysical features

<u>*Distance from the channel pore axis*</u>

The distance of mutation site from channel pore axis was an important biophysical feature that help identify functionality vital regions in the KCNQ1 Protein. The figure S1 shows two different mutation sites with blue color site in VSD region and red color site in PD region.

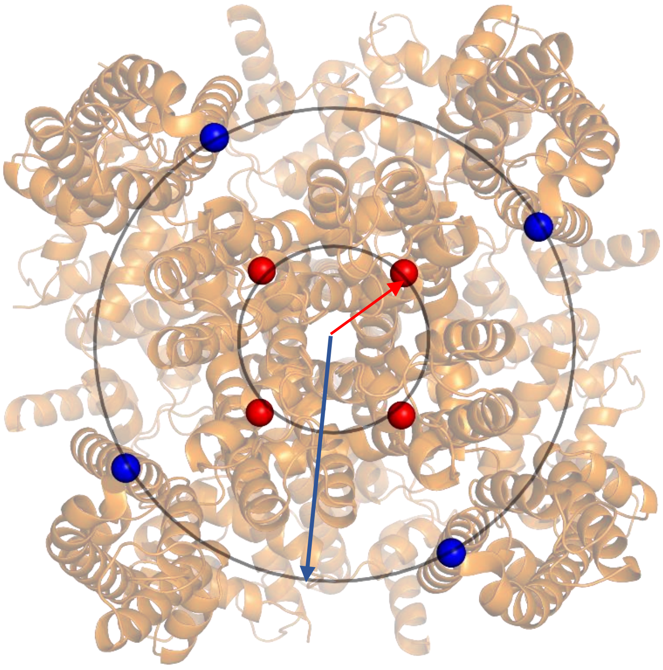

Figure S1: Distance from the channel pore axis

*Burial of mutation site in the membrane*

The depth of the mutation site relative to membrane thickness was weighted based on three regions defined as mutation on the membrane, mutation at the protein solution interface, and mutation outside the membrane. The thickness of the membrane was considered as 31.4 Å. A decaying cosine function was used at the interface for a smooth transient from the site being on the membrane to outside the membrane as shown in Figure S2. The criteria for depth of mutation are as follows:

$$Depth\big(Distance(z)\big) = \begin{cases} 1, & z < 11.25\ \text{Å} \\ \dfrac{1+\cos\left(\pi \times \frac{z-11.25}{23.75-11.25}\right)}{2}, & 11.25\ \text{Å} \le z < 23.75\ \text{Å} \\ 0, & z > 23.75\text{Å} \end{cases} \quad \ldots\ldots\ldots(1)$$

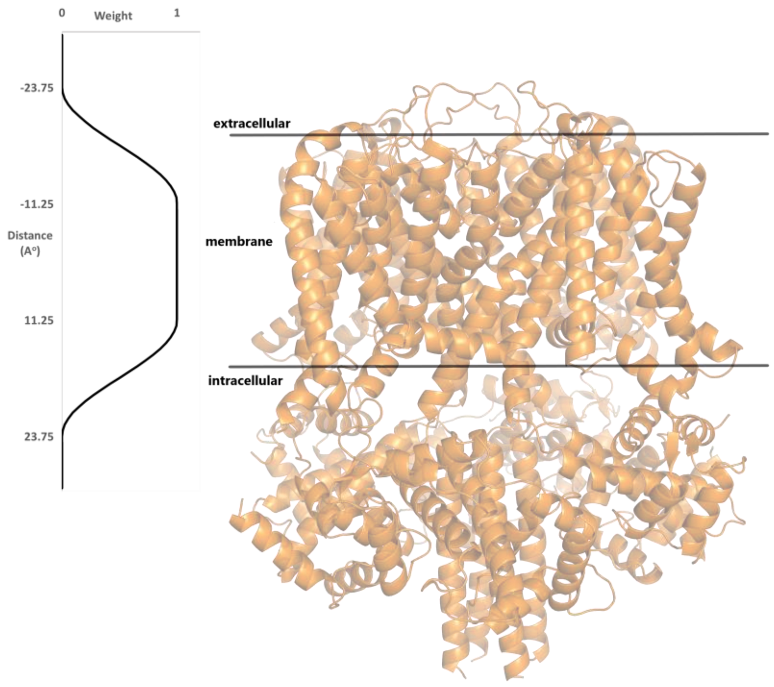


Figure S2: Depth of the site of mutation on the membrane

*Hydrophobicity an indicator of free energy change at the mutation site*

Hydrophobicity plays a vital role in the folding, structural stability, and functioning of the membrane protein. We utilized a previously developed hydrophobic scale for mammalian alpha helical membrane protein by Koehler *et al* (1). Their work distinguishes amino acid preference for solution, interface, and membrane region as shown in Figure S3. For consistency, we incorporated the same definition and thickness of interface and membrane. The value of hydrophobicity for an amino acid was taken based on the region where the site of the mutation exists.

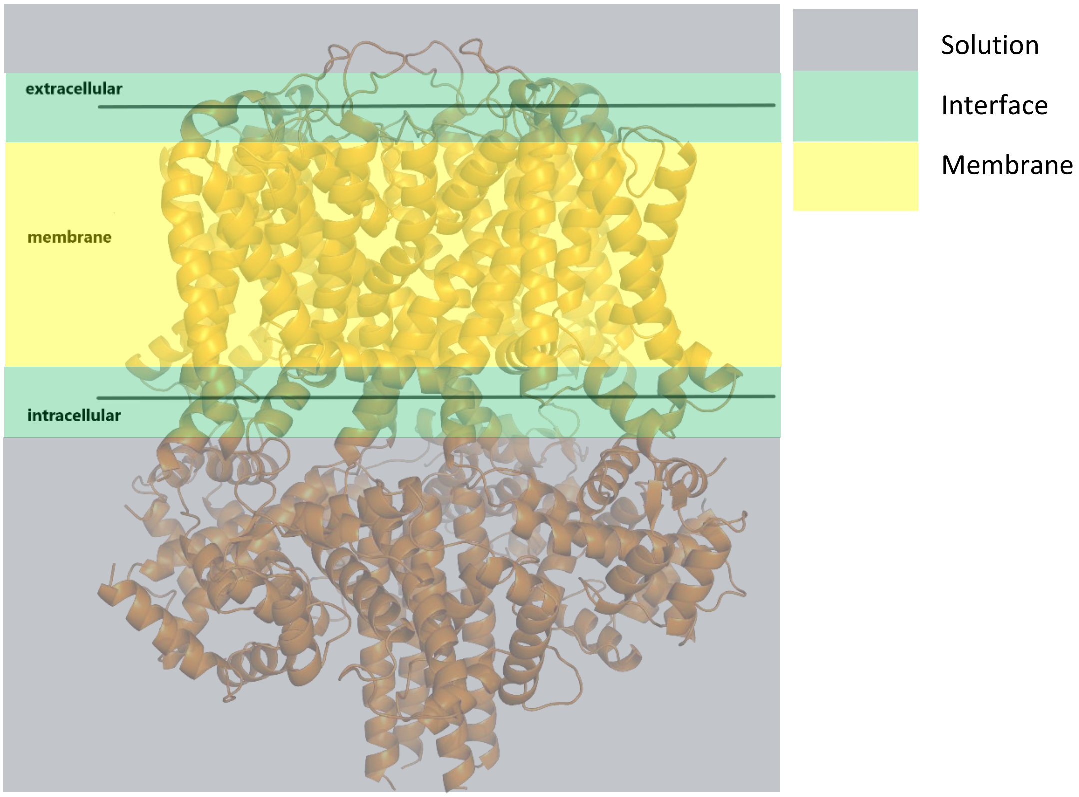


Figure S3: Definition of three regions of hydrophobicity utilized in this work

*Functional density for polarizability and hydrophobicity*

In addition to structure-based feature discussed previously, we computed an average physiochemical property of the neighborhood around the site of mutation. The concept of functional density from Kroncke *et al* (2) quantifies the functional critical spots within the protein structure. This method is based on k-nearest neighbors' algorithm, wherein average physiochemical property (for this study were polarizability and hydrophobicity) around the site of mutation was weighted by inverse of their distance from the site of mutation. For instance, functional density for polarizability is calculated as follows:

$$f(x,i) = \frac{\overline{Polarizability} + \sum_{i}^{Variants} Weight(Distance(x,i) \times Polarizability(i))}{1 + \sum_{i}^{Variants} Weight(Distance(x,i))} \quad \ldots.\ldots (2)$$

$$Weight(Distance(x,i)) = \begin{cases} 1 & Distance < X_{Start} \\ \dfrac{1 + \cos\left(\pi \times \dfrac{Distance(x,i) - X_{Start}}{X_{end} - X_{Start}}\right)}{2} & X_{Start} \leq Distance < X_{End} \\ 0 & Distance \geq X_{End} \end{cases} \quad .(3)$$

where $f(x,i)$ is the functional density at the site of interest with i$^{th}$ neighbor residue within the threshold distance and to take prior knowledge into account, the average polarizability from the dataset was taken as pseudo count represented as $\overline{Polarizablity}$ in the equation 2. The summation runs for all the neighboring residue existing within the threshold distance from the residue of interest. A decaying cosine function of thickness 0.5 Å was used to have a smooth transition of weight towards the end of the neighbor threshold. For polarizability, we used two neighborhoods with different radius ($X_{End}$) with first shell of contacting residues at 6.5 Å and a second shell at 12 Å (see Figure S4). We used similar implementation for hydrophobicity around the site of mutation except those two neighborhoods with different radius ($X_{End}$) with first shell of contacting residues at 1 Å and a second shell at 6.5 Å. These radial distances are based on distance from $C_ß$ of native amino acid.

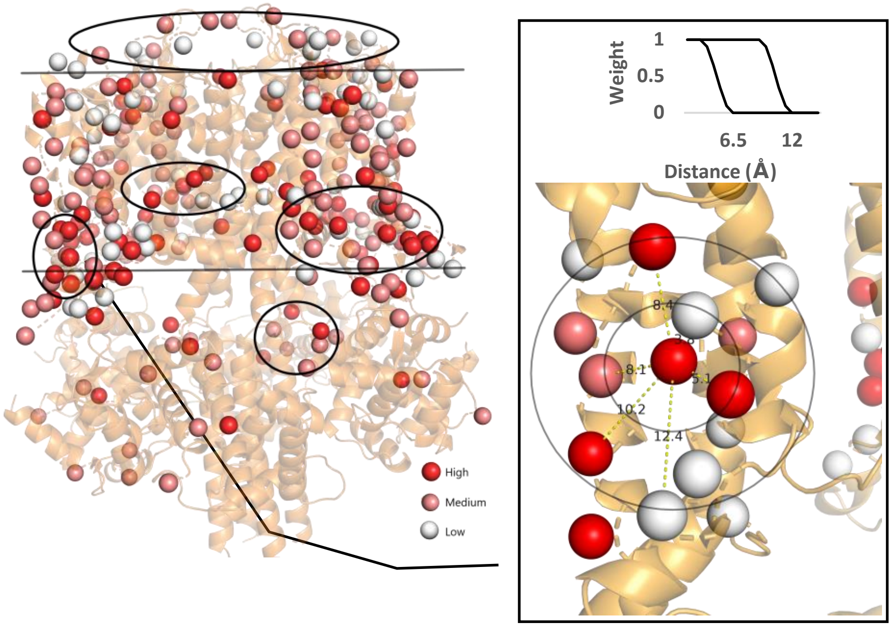


Figure S4: This figure depicts polarizability distribution with clusters of high, medium, and low polarizability. This also captures the concept of functional density by quantifying these clusters of polarizabilities for different neighborhood size.

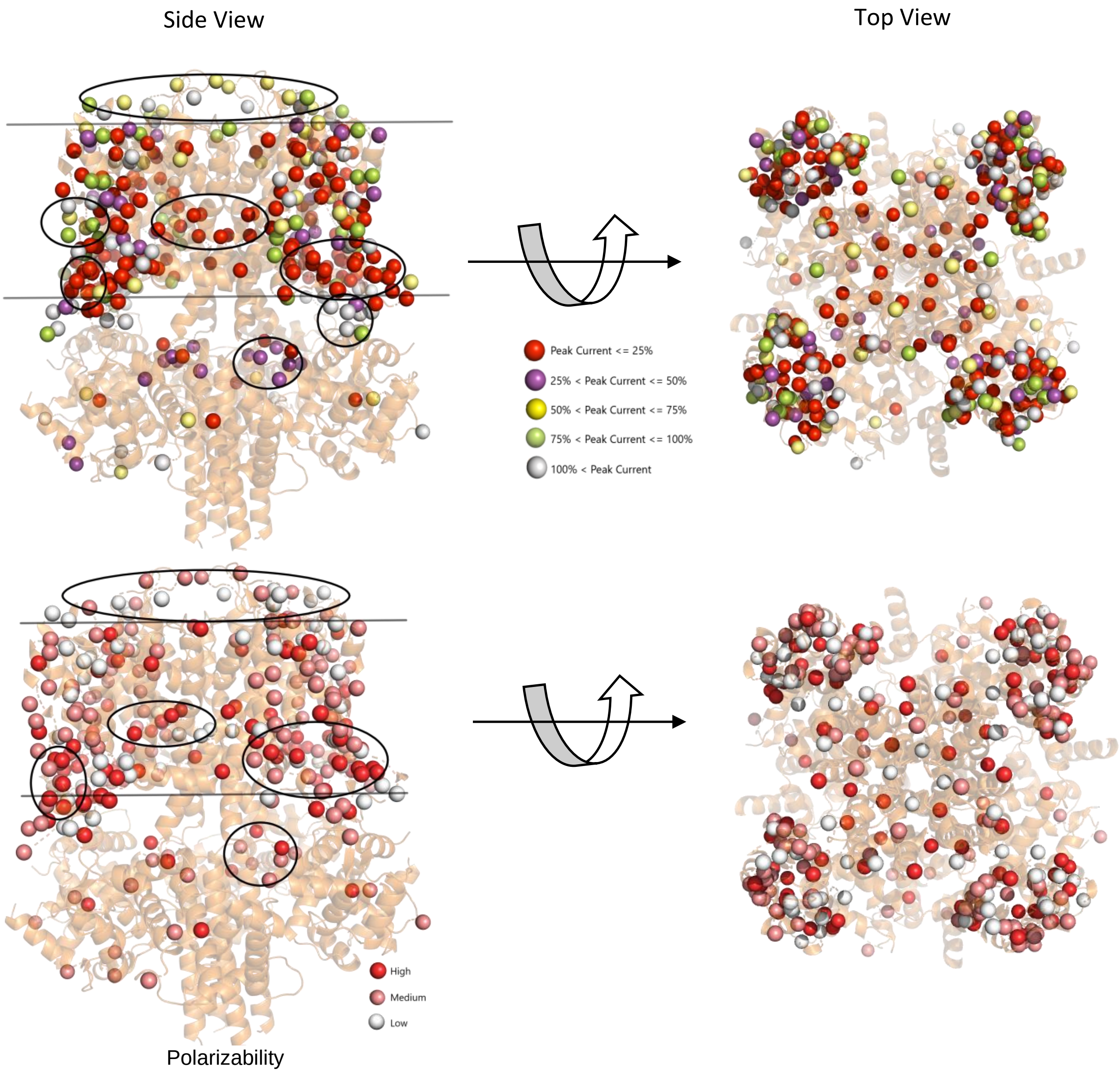


Figure S5: Correlation of peak current with polarizability at different pockets in the protein structure

*Neighbor vector captures buried site in the protein structure*

The sites that are buried within the core of the protein are highly conserved, critical to the stability, and functioning of the protein. Mutation at the conversed sites (or less exposed) can immensely impact the stability and functioning of the protein. Therefore, to identify these important sites from the protein structure, we recall neighbor vector definition from Durham *et al* (3). Neighbor vector considers the spatial orientation of neighboring amino acids within a radius of 11.4 Å from the $C_{ß}$ of the native amino acid. The magnitude of neighbor vector implies exposure of residue of interest in the protein environment wherein long lengths of this vector $\cong 1$ signify high exposure or otherwise.

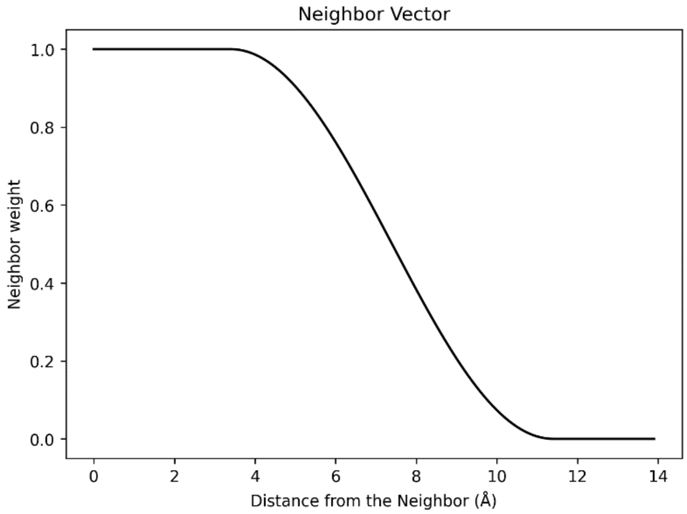


Figure S6: A definition of neighbor that includes a smooth transition function used in the neighbor vector algorithm with lower bound at 3.3 Å and upper lower at 11.4 Å

*Amino acid parameters and physicochemical properties*

In addition to structure-based and environment-based features, we also introduce few physiochemical parameters of amino acid that improves the prediction. There properties were steric parameter, normalized vanderWaal volume, polarizability, number of hydrogen donor, and number of hydrogen acceptor sites (4). These parameters are list in the Table 1 for 20 amino acids.

The steric parameter quantifies the complexity, symmetry, and branching at the $C_\alpha$ from the structure of amino acids(4). The polarizability is related to molar refractivity as

$$\alpha = \frac{3}{4\pi N} \cdot \frac{M}{d} \cdot \frac{n^2 - 1}{n^2 + 2}$$

Where n is index of refraction, M is molecular weight , d is density, and N is number of atoms(4).

The normalized vanderWaal volume is defined as

$$v(side\ chain) = \frac{V(side\ chain) - V(H)}{V(CH_2)}$$

This normalized volume is thus, 0 for glycine and 1 for alanine(4).

Table S1: Amino acid parameters

| AMINO ACIDS | STERIC PARAMETER | POLARIZABILITY | VOLUME | NO. OF DONOR SITES | NO. OF ACCEPTOR SITES |
|---|---|---|---|---|---|
| **ALA** | 1.28 | 0.05 | 1.00 | 0 | 0 |
| **GLY** | 0.00 | 0.00 | 0.00 | 0 | 0 |
| **VAL** | 3.67 | 0.14 | 3.00 | 0 | 0 |
| **LEU** | 2.59 | 0.19 | 4.00 | 0 | 0 |
| **ILE** | 4.19 | 0.19 | 4.00 | 0 | 0 |
| **PHE** | 2.94 | 0.29 | 5.89 | 0 | 0 |
| **TYR** | 2.94 | 0.30 | 6.47 | 1 | 1 |
| **TRP** | 3.21 | 0.41 | 8.08 | 1 | 0 |
| **THR** | 3.03 | 0.11 | 2.60 | 1 | 1 |
| **SER** | 1.31 | 0.06 | 1.60 | 1 | 1 |
| **ARG** | 2.34 | 0.29 | 6.13 | 3 | 0 |
| **LYS** | 1.89 | 0.22 | 4.77 | 3 | 0 |
| **HIS** | 2.99 | 0.23 | 4.66 | 2 | 2 |
| **ASP** | 1.60 | 0.11 | 2.78 | 0 | 2 |
| **GLU** | 1.56 | 0.15 | 3.78 | 0 | 0 |
| **ASN** | 1.60 | 0.13 | 2.95 | 1 | 0 |
| **GLN** | 1.56 | 0.18 | 3.95 | 1 | 1 |
| **MET** | 2.35 | 0.22 | 4.43 | 0 | 0 |
| **PRO** | 2.67 | 0.00 | 2.72 | 0 | 0 |
| **CYS** | 1.77 | 0.13 | 2.43 | 0 | 0 |

**Evolutionary features**

We used PSI blast technique to characterize the likelihood of an amino acid substitution from the perspective of protein evolution(5). This technique was implemented by obtaining position specific scoring matrix (PSSM) by searching though uniref50 databases(6) and NCBI non-redundant sequence databases(7) with PSI-BLAST for four iterations. The E-value inclusion threshold was set to 0.00001. A PSSM matrix of size 676 x 20 gives log ratios of frequency of 20 amino acids to occur at 676 sites for KCNQ1 protein relative to the frequency of wild type amino acid.

$$Likehood\ of\ Amino\ acid\ substitution = \lambda\, ln \frac{P_A}{P_A^o}$$

where $P_A$ is the probability of amino acid A at a position, $P^o{}_A$ is the expected probability for wild type amino acid, and λ is a scaling factor built in PSI-BLAST(5).

The difference of PSSM score between mutant amino acid (A) and native amino acid (B) from two databases are as follows:

$$PSSM(NR) = \lambda\left(ln\frac{P_{A_{NR}}}{P_{A_{NR}}^o} - ln\frac{P_{B_{NR}}}{P_{B_{NR}}^o}\right)$$

$$PSSM(uniref50) = \lambda\left(ln\frac{P_{A_{uniref50}}}{P_{A_{uniref50}}^o} - ln\frac{P_{B_{uniref50}}}{P_{B_{uniref50}}^o}\right)$$

PSSM(NR) and PSSM (uniref50) were evolutionary features used in this work that measure the perturbation due to amino acid substitution. Higher the perturbation, more likely it is to have a functional or structural impact on the protein due to mutation.

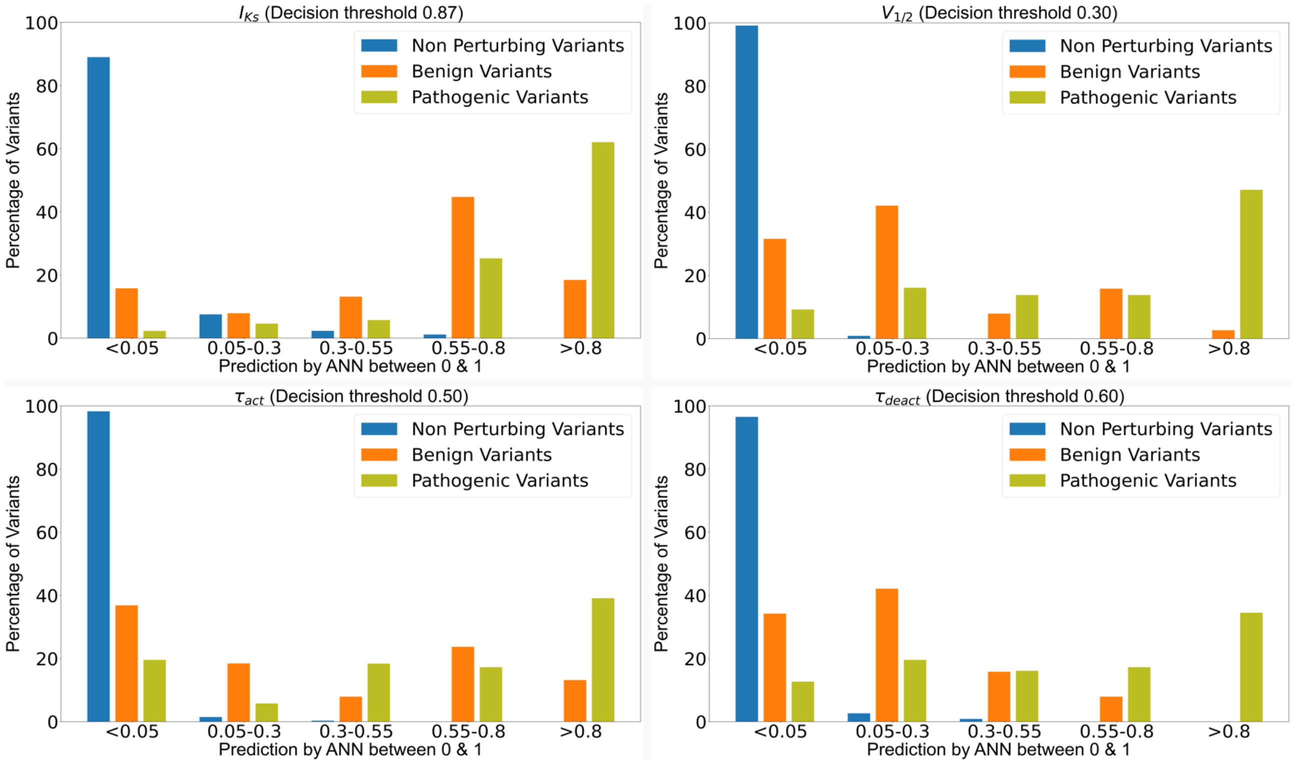


Figure S6: Distribution of Prediction by ANN for non-perturbing, benign, and pathogenic variants depicting that ANN can distinguish these variants by predicting in three different regions between 0 and 1. Decision threshold is between benign and pathogenic variants

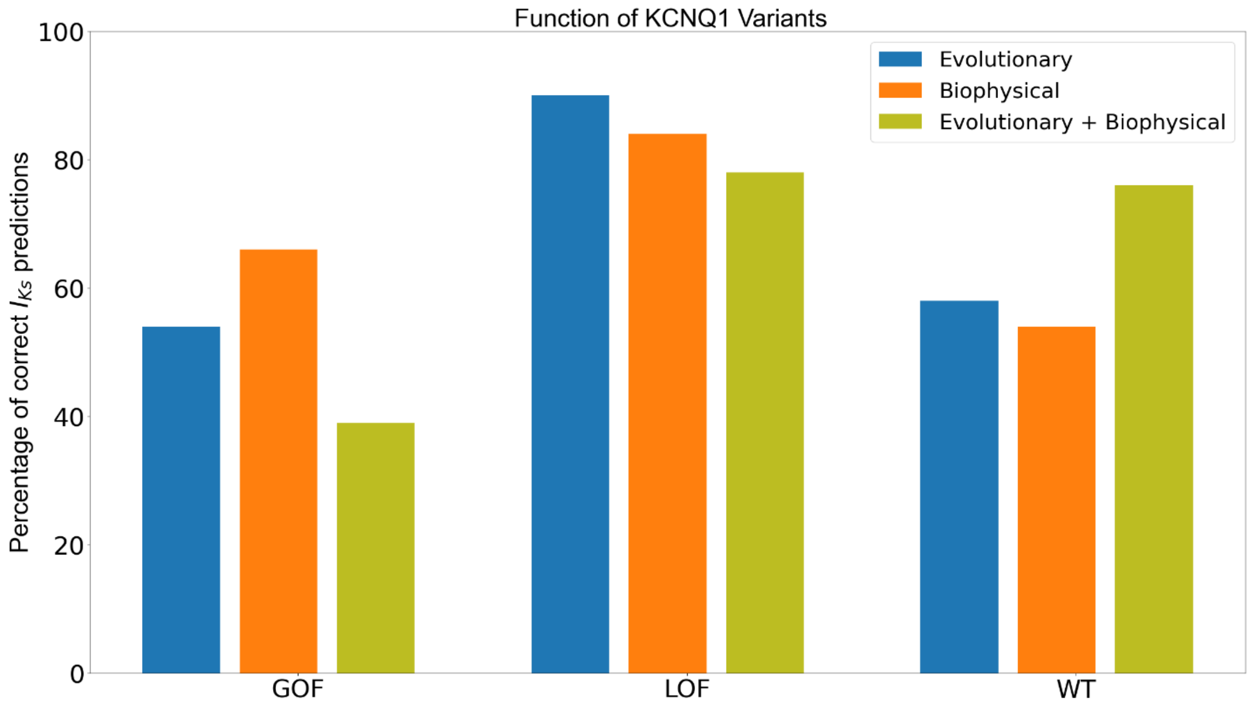


Figure S7: Percentage of accurate predictions for GOF, LOF and WT-like based on peak current density by the three ANNs models considered in this study

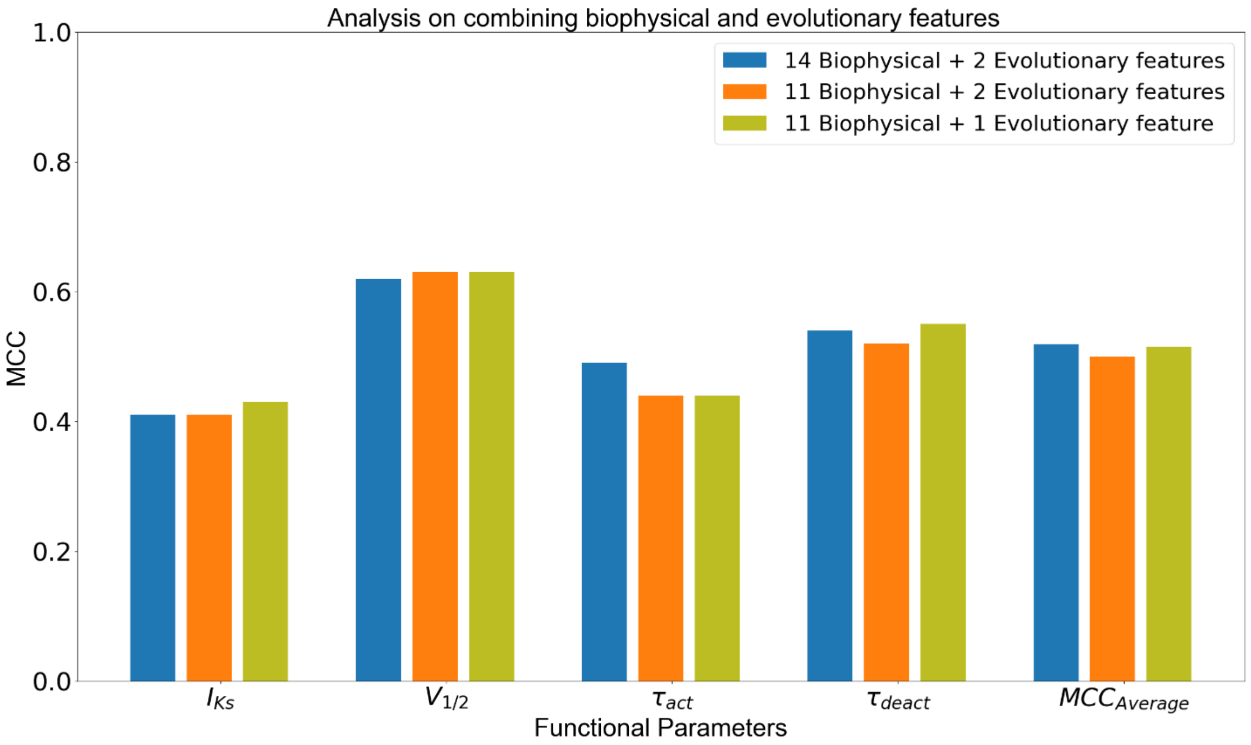


Figure S8: Exclusion of 3 biophysical features and 1 evolutionary feature does not affect the performance when evolutionary and biophysical features are combined
